# From connectome to effectome: learning the causal interaction map of the fly brain

**DOI:** 10.1101/2023.10.31.564922

**Authors:** Dean A. Pospisil, Max J. Aragon, Sven Dorkenwald, Arie Matsliah, Amy R. Sterling, Philipp Schlegel, Szi-chieh Yu, Claire E. McKellar, Marta Costa, Katharina Eichler, Gregory S.X.E. Jefferis, Mala Murthy, Jonathan W. Pillow

## Abstract

A long-standing goal of neuroscience is to obtain a causal model of the nervous system. This would allow neuroscientists to explain animal behavior in terms of the dynamic interactions between neurons. The recently reported whole-brain fly connectome [1–7] specifies the synaptic paths by which neurons can affect each other but not whether, or how, they do affect each other in vivo. To overcome this limitation, we introduce a novel combined experimental and statistical strategy for efficiently learning a causal model of the fly brain, which we refer to as the “effectome”. Specifically, we propose an estimator for a dynamical systems model of the fly brain that uses stochastic optogenetic perturbation data to accurately estimate causal effects and the connectome as a prior to drastically improve estimation efficiency. We then analyze the connectome to propose circuits that have the greatest total effect on the dynamics of the fly nervous system. We discover that, fortunately, the dominant circuits significantly involve only relatively small populations of neurons—thus imaging, stimulation, and neuronal identification are feasible. Intriguingly, we find that this approach also re-discovers known circuits and generates testable hypotheses about their dynamics. Overall, our analyses of the connectome provide evidence that global dynamics of the fly brain are generated by a large collection of small and often anatomically localized circuits operating, largely, independently of each other. This in turn implies that a causal model of a brain, a principal goal of systems neuroscience, can be feasibly obtained in the fly.

## Introduction

A fundamental barrier to resolving a causal model of the nervous system is that causal relationships in the brain cannot be inferred solely from passive measurements of neural activity [8, 9]. Direct perturbation of neural activity (e.g., optogenetic stimulation) confronts this problem and, therefore, has been an area of intense methodological research and resulting progress. Yet a clear approach for how to employ these tools to obtain a causal model of neural activity has not emerged, Here we introduce a novel combined statistical and experimental strategy, and demonstrate that it can efficiently learn a causal model of the fly brain.

We adapt a technique known in the statistical literature as ‘instrumental variables’ (IV) [10]. This technique was developed to estimate causal relationships in observational data. It relies critically on the stringent requirements of an IV: it only directly affects an observed variable that putatively effects the outcome of interest—absent that effect, the IV is independent of all variables. The observed relationship between the IV and outcome is then strictly a result of the causal effect of interest. Yet, despite the stringency of these requirements, optogenetic stimulation plausibly meets them [11, 12]. Optogenetic stimulation affects neural activity, it is independent of neural activity because it is controlled by the experimenter, and it only acts on the brain through neurons that have expressed opsins. Thus the IV approach could in principle be used to estimate causal effects between neurons.

However, there are two fundamental problems with this approach applied to an entire nervous system. First, naively estimating parameters between every pair of neurons in the fly (*∼* (10^5^)^2^) would require intractable amounts of data. This problem in particular is an even more insurmountable barrier to learning causal models of higher order organisms with far more neurons (e.g. for mice *∼* (10^8^)^2^ potential effects [13]). Second, it would be infeasible in the fly—and most organisms—to independently stimulate and record from all neurons at once. The effectome thus would need to be gradually constrained across many experiments. It is unclear how to order experiments such that insight into whole brain dynamics are achieved efficiently. Here we show that the Flywire connectome [1–7] provides a feasible path to surmounting both of these barriers in the fly.

First, a principled approach to improving the data efficiency of an estimator is to place priors on the parameters being estimated. The connectome can be used as a prior on effects between neurons in the fly brain: distant neurons with no synaptic contacts are unlikely to directly affect each other. The connectome of the fly is, fortuitously, exceedingly sparse (*∼* 0.01% of neuron pairs form a synaptic contact [14]): thus a strong prior can be placed on the vast majority of the effectome. Furthermore, with synapse-level neurotransmitter predictions [7], priors can be placed on the sign of effects between the neurons that are connected (the confidence of those predictions can naturally be incorporated into the strength of the prior).

Second, because the Flywire connectome is a complete connectivity map of the fly brain, it is uniquely suited to guide efficient estimation of the effectome. Neuroscientists tend to study a given neural circuit because it has previously been studied. A data-driven approach to proposing neural circuits of interest could reveal important computations that had not yet been considered. Ideally such a method would discover independent neural circuits with interpretable dynamics and rank them by their total effect on the brain. In order to predict which neural interactions contribute most strongly to whole-brain dynamics, we analyze the eigenmode decomposition of the connectome. The eigenvectors of the connectome, ordered by the magnitude of their eigenvalues, provide sets of neurons, and patterns of neural activity therein, that under our ‘connectome prior’ are predicted to have the greatest total effect on the brain. The dynamics of these circuits can be described with only two parameters (the real and complex part of their associated eigenvalue) and are therefore interpretable. We thus provide an experimentally tractable paradigm for learning a complete causal model of the fly central nervous system that is uniquely enabled by the properties of the whole brain connectome. Specifically, we show that the sparse connectivity between neurons drastically improves the efficency of estimation of causal effects and our discovery that small populations of neurons underlay dominant dynamical modes suggests whole brain dynamics can be constrained with a series of highly targeted, and thus feasible, experiments.

## Results

Here we first lay out an experimental setup and associated statistical model of neural activity in the fly. Next, we outline how to use optogenetic perturbation as an instrumental variable to infer the direct causal effects between neurons in the context of a linear dynamical system. We then show how in the presence of confounds the classic regression estimator will return biased results whereas the IV approach gives a consistent estimator. By simulating the entire fly connectome, we demonstrate that the IV approach provides consistent estimates of ground truth neural connectivity. We find that because there exists a massive number of potential downstream neurons from any given neuron, the IV regression estimator converges very slowly. This motivates using the connectome as a prior on the IV weights so that the estimator remains consistent yet orders of magnitude more efficient. Finally, we analyze our ‘connectome prior’ to reveal thousands of proposed circuits ranked by their predicted total effect on the brain. We analyze two of these circuits, and find that one recapitulates a proposed circuit for computing opponent motion and the other provides a dynamical mechanism for visual spatial selectivity.

Our motivating setting is a typical optogenetic experiment in the fly (Fig 1): a population of neurons’ activity is observed (target neurons, within microscope FOV, black oval), a subset of these observed neurons expresses opsins (source neurons, under red laser light), these are driven by *n_l_* independent stimulation channels where the modulation signal is IID across time (red trace over laser), and the remaining neurons remain unobserved—these can also be unobserved non-neuronal processes recurrently interacting with the observed neurons (grey circles outside of FOV).

**Fig. 1:**
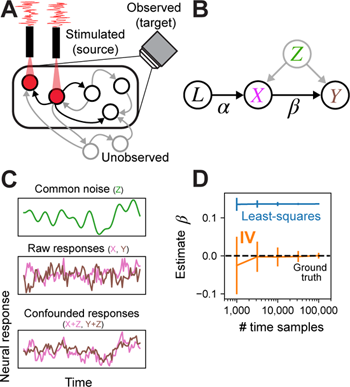
Schematic of experimental setting, model, and demonstration of instrumental variables in simulation. **(A)** Example of single fly experiment. Some sets of neurons in the fly are observed (black circles within FOV of camera), some subset of these express opsin and are being stimulated (neurons under red light) with a white noise pattern (red trace). Another set of neurons may be unobserved. All neurons can have directed interactions via synapses (arrows), only the effects of stimulated neurons can be estimated (stimulated neurons arrows to observed neurons black). **(B)** In this setup, the laser can be cast as an instrumental variable that directly affects only the opsin-expressing neurons (arrow labeled *α* from *L* to *X*). Stimulated neurons in turn have a direct effect on downstream observed neurons (arrow *β* from *X* to *Y*), but common unobserved inputs may corrupt attempts to estimate the direct relationship. The IV approach uses the joint relationship between the laser and downstream neurons to determine the direct effect of the stimulated neurons on postsynaptic neurons. **(C)** Simulated example of confounding effect *Z*, which is given by a slow drifting signal (smooth green trace, top). Raw uncorrupted responses of neurons *X* and *Y* are independent (pink and brown traces, middle), meaning there is no connection from *X* to *Y*. Observations of *X* and *Y* after adding the confounding signal *Z*, resulting in substantial correlation (bottom). **(D)** The least-squares estimate of the weight from *X* to *Y* (blue; mean *±*1SD) exhibits large bias regardless of sample size, whereas IV estimate (orange) converges to the true effectome weight of zero (black dashed).

The graphical model we associate with this setup identifies the lasers as the instrumental variable that uniquely drives the source population (*X*) via a linear transformation (*α*), the source population drives the target population (*Y*) via *β* (the causal estimand), and both receive common inputs from unobserved confounders *Z*(Fig 1B). The fundamental difficulty of fitting neural models to observed data is that even if *β* = 0, *Z* can induce spurious dependence between the source and target population. By casting the laser as an instrumental variable, we avoid an intractable source of confounds when limited to observational data, namely confounds via unobserved common inputs. Below we provide toy simulations demonstrating how the naive regression approach fails but IV succeeds in this scenario.

### Instrumental variables approach is not confounded by unobserved variables

The principal challenge to inferring causal effects solely through passive observation of neural activity is the possibility of unobserved confounding inputs. Potential unobserved confounders in the fly are numerous. Typically a small subset of the nervous system is imaged at a time: thus input from unobserved neurons could confound the observed neurons. Whole-brain imaging is possible in the fly [15–17], but resolving single neuron dynamics remains a challenge due to the density of neuropil. Techniques like constrained non-negative matrix factorization can segment activity profiles [18] but their accuracy in extracting single-neuron dynamics remains untested. Even if whole brain activity could be recorded accurately, afferent activity from the peripheral nervous system and ventral nerve cord could likely be a source of common variability. To date, there are no imaging methods that can resolve single-neuron activity throughout the whole body. Furthermore, even if every neuron’s activity were observed, diverse neuromodulatory processes, like hormones and neuropeptides, cannot be inferred from neural data alone and could induce common modulatory effects over various timescales [19–21]. Finally, measurement error, including physiological artifacts like brain movement [22], can act as a source of common variability. Collectively, there is a high probability of unknown confounding variables preventing valid causal inference during passive observation of neural activity in the fly brain.

A simple case that demonstrates the effect of confounding variables is one in which two neurons, *X* and *Y*, have no causal effect on each other (*β* = 0 between *X* and *Y*, Fig 1B) but there is a common unobserved input to both from *Z* (e.g, an unobserved neuron). We simulate this common input as a slow drift (Fig 1C, top panel) that is added to both the raw independent responses of *X* and *Y* (pink and brown trace Fig 1C, middle panel), creating slow common fluctuations in both (Fig 1C, bottom panel). Even though there is no causal effect of X on Y (Fig 1D ground truth weight is 0), the least-squares estimate incorrectly converges on a positive weight, whereas IV converges on 0 because there is no correlation between the laser and *Y*. Thus IV is not corrupted by unknown, unobserved inputs.

### Instrumental variables approach accurately estimates fly effectome

To demonstrate the estimator can, in principle, estimate the fly effectome, we applied it to a simulation of the entire fly brain during stimulation of a single source neuron and a whole brain recording (see Methods). We set our ground truth causal effect matrix entries to be proportional to the number of synapses multiplied by the ‘sign’ of those synapses—whether they were inhibitory or excitatory (Fig 2A). We divided the matrix by the magnitude of its first eigenvalue so that its activity was stable. We then applied the raw IV estimator to the simulated data. We found that the estimator on average was accurate (Fig 2B, scatter centered around diagonal) but the bulk of variability in the estimator came from estimates of weights whose true value was zero. This was not because the estimator gave noisier estimates of weights that are 0, but because the vast majority of weights are 0 due to the sparsity of the fly connectome (0.01% of neuron pairs make synaptic contact). Non-zero estimates of these weights were thus the dominant contribution to total estimation error.

**Fig. 2:**
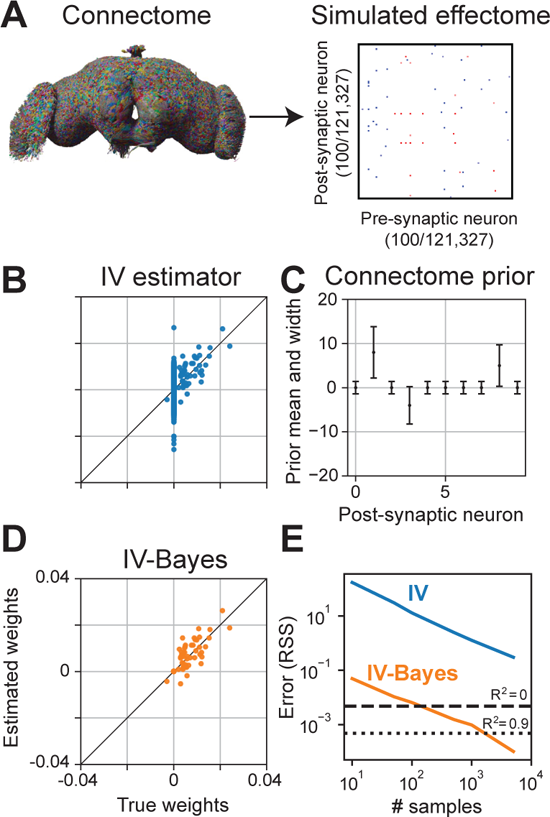
Inferring the effectome using standard and Bayesian approaches on simulated data. **(A)** We used the connectome to set the effectome weights for a whole-brain simulation using all 121,327 neurons. Synaptic weights were set proportional to synapse count, with positive (negative) sign for excitatory (inhibitory) synapses. **(B)** Instrumental variable (IV) estimates of post-synaptic weights of a single example neuron. The majority of the error falls along the vertical line where true weight equals 0 (which is the majority of the weights, due to the sparsity of the fly connectome). **(C)** Mean *±*2 SD of an independent Gaussian “connectome prior” on each weight in the effectome. We set the prior mean to be proportional to the signed synapse count of the connectome, and variance equal to the absolute value of the mean plus a small constant, so that the prior width is non-zero between neurons with no known synapses. **(D)** IV-Bayes estimator shows less variance than raw IV (B). **(E)** Error of estimated weights decreases with the number of samples for both estimators, but IV-Bayes gives several orders of magnitude faster convergence.

The high variability of our estimator reflects a fundamental problem in fitting a model of the entire brain: the number of parameters is large. Yet because the fly connectome is available and, critically, it happens to be highly sparse, we can dramatically increase the efficiency of our model estimate by making the assumption that neurons with no synaptic contacts are unlikely to directly affect each other. We do so in a principled manner that maintains the consistency of the IV estimator by extending it to the Bayesian setting. Specifically, we put priors on the estimated synaptic weights between neurons according to the fly connectome.

### Using the fly connectome as a prior on the effectome

Motivated by the high variability of the IV estimates, we took a Bayesian approach to reduce error by placing a prior on the model weights. We refer to the resulting estimator as IV-Bayes. The prior distribution on the weights was Gaussian with mean proportional to the product of the synaptic count and sign (positive for excitatory synapses, negative for inhibitory synapses) of the connectome (Fig 2C). The variance of the prior was proportional to the absolute value of the mean, plus a small constant. This shrinks the effectome weights between non-anatomically connected neurons to be close to zero, and the weights of anatomically connected neurons toward the synapse count (scaled by a global constant), while allowing them to be zero if warranted by the data. This could be the case if the experimental conditions never lead to the simultaneous activation of those two neurons thus no effect. We add a small constant to the prior variance so that if a synapse exists where none was found in the original connectome, the estimator will, with enough data, still identify the correct causal effect.

We evaluated the IV-Bayes estimator using a simulated dataset generated with effectome weights set to a corrupted version of the ground truth connectome weights, thus creating a mismatch between the effectome weights and the prior mean (see Methods). This corruption could reflect natural variation between flies’ connectomes, measurement error in the connectome, or variation in synaptic transmission efficacy between neurons. We found that the IV-Bayes estimates out-performed the standard IV estimator (Fig 2D). In particular, the high variability of the IV estimate for zero weights was quenched in the Bayes-IV estimate. Intuitively, as long as the connectome on average provides information about the strength of causal interactions between neurons, it should out-perform standard IV.

To quantify the relative efficiency of the naive IV approach and IV-Bayes, we sampled effectome matrices as described above and then evaluated the average residual sum of squares (RSS) of the two estimates as a function of the number of samples (e.g., duration of experiment). As expected we found that both estimators’ RSS decreased with increasing samples (Fig 2E, blue and orange trace slope downwards). Yet we found the RSS of the IV estimator is at least an order of magnitude higher than that of the IV-Bayes estimator across number of samples (blue above orange). In terms of fraction variance explained IV-Bayes explains the vast majority of variance for the maximal number of time samples (orange trace below dotted line on right) but the raw IV estimator is still too noisy to achieve a positive quantity of fraction variance explained (blue trace above dashed line). Thus in simulation IV-Bayes provides at least an order of magnitude improvement in converging to the ground truth causal effects.

### Road map to the effectome

We now turn to a strategy of learning the entire fly effectome using Bayes-IV. Our goal is to learn the entire effectome using genetically identical flies under the same experimental conditions (e.g., during walking, courtship, or resting). For example, making estimates of the effectome only during walking would enable us to infer which neurons on average directly affect each other during the walking state.

Our simulations thus far have focused on the stimulation of a single neuron while the entire fly brain is observed. A conceptually straight-forward approach to learning the entire effectome would be to stimulate one of each of the 121,327 fly neurons in 121,327 different flies while imaging the whole brain. This amounts to learning one column of the causal effect matrix per a fly (Fig 3A). Yet, amongst other technical issues with this approach, performing this number of experiments would be difficult. Learning the effectome more rapidly requires a higher rank perturbation. In the case of optogenetics, the rank could be increased with 2-photon holographic stimulation so that arbitrary sets of neurons could be driven independently [23]. If every neuron in the fly brain was stimulated independently while every neuron was observed, the entire fly effectome could be identified within a single experiment (Fig 3B).

**Fig. 3:**
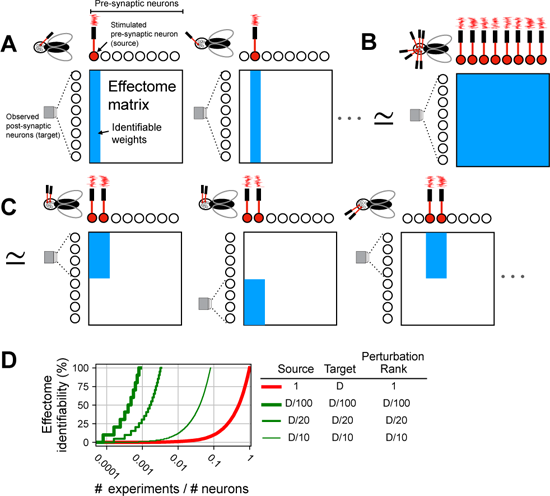
Schematic of perturbation experiment types. **(A)** (left) Single fly experiment with whole brain observation (microscope FOV encompasses all post-synaptic neurons), single neuron stimulation (laser drives filled red neuron), and total rank of stimulation is 1 (only one laser). As a result all post-synaptic weights from the stimulated neuron are identifiable (blue column of weight matrix filled). (middle) To learn all other effectome weights in this setup would require as many flies as neurons, as each individual neuron is stimulated. **(B)** Single fly experiment in which all weights are identifiable: whole brain observation, whole brain stimulation, and rank of stimulation is equal to the number of neurons in the brain (same number of lasers as neurons). **(C)** (left) Single fly experiment with partial brain observation (FOV encompasses half of neurons), partial brain stimulation (two neurons red filled), and rank of stimulation is equal to that of the source (2). (middle) To identify all weights of source neurons the experiment is repeated in another fly but with different target neurons, all effectome weights can be identified piecemeal in this manner. **(D)** Convergence rate to full identifiability of the fly effectome as a function of the fraction of number of experiments over the total number of neurons. Different traces reflect different experimental settings. Columns in legend are three primary ways experiments can vary. Source is the number of neurons that are being stimulated. Target is the number of neurons observed, we assume the source neurons are also observed. Perturbation rank is the dimensionality of the perturbation method. *D* is the total number of neurons.

A common issue with the two hypothetical approaches described above is the difficulty of observing all neurons simultaneously. Resolving single neurons from densely labeled neuropil recordings is an open problem. Similarly, independently targeting all neurons for whole-brain patterned stimulation is infeasible given the constraints of diffraction-limited optics. Moreover, the amount of energy required to optogenetically target all neurons in the brain would likely heat the brain tissue and interfere with its function. Here, we propose an alternative approach to learning the effectome that could be achieved in a feasible number of experiments, permits accurate identification of neurons, and leverages existing technologies.

We propose that sparse populations of neurons should express both opsin and GCaMP so that they can be both stimulated and observed simultaneously. Holographic optogenetic stimulation can then be used to drive each neuron independently. Recently developed experimental techniques allow recording from the fly brain for up to 12 hours [24]: this would allow for a block of the effectome weights for a complete subset of neurons to be learned over the course of a single experiment (Fig 3C, see Appendix B for further detail).

The efficiency of this approach depends on the size of the population that expresses both opsin and GCaMP (source population). To evaluate efficiency, we consider percent identifiability as a function of number of experiments per neurons total. Percent identifiability is simply the percent of the effectome that could be learned given an unlimited amount of data (i.e., consistency for a portion of of the effectome matrix). We consider this in units of experiments per neuron because for a given application the investigator may want to learn the effectome of a sub-population of neurons.

For reference, we first considered the exhaustive approach of stimulating each neuron individually (Fig 3A). In this case, the effectome would accumulate slowly and require the same number of experiments as neurons (Fig 3D thick red trace intersects 100 % at 1). In general with a perturbation rank of 1, at least as many experiments as neurons would be needed. On the other hand, our sparse patterned stimulation approach achieves full identifiability with orders of magnitude greater efficiency, even if only 5 % of neurons are observed for each experiment (medium green trace intersects 100% at 420 experiments). This rate could be achieved even more rapidly with higher expression rates (thicker green trace) but at the cost of increased difficulty of identifying and stimulating larger populations of neurons. Thus with existing technologies the entire effectome could be learned in a feasible number of experiments.

Although our sparse stimulation and recording strategy provides a straight-forward approach for efficiently learning the effectome, we have not specified a sequence in which these experiments should be conducted. If stimulation were performed randomly, a large fraction of the effectome would likely need to be estimated before meaningful circuitry was uncovered. We now describe a strategy for systematically choosing subsets of neurons that will, with high likelihood, provide valuable scientific insight.

### Identifying dominant circuits using the connectome

Here we demonstrate a data-driven method for ranking sets of source neurons most likely to form circuits with a large effect on the fly nervous system. Specifically, we consider a recurrent neural network model of whole-brain activity given by

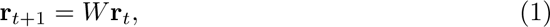

where **r***_t_* is a vector denoting the activity of all neurons at time *t* and *W* is the effectome weight matrix, which (for these analyses) we set to the scaled, signed synaptic counts extracted from the connectome. To analyze the fly brain’s dynamical properties, we perform an eigendecomposition of the weight matrix *W*, which decomposes global dynamics into patterns of neural activation called eigenvectors with simple two-dimensional rotational dynamics determined by an associated eigenvalue. Below, on the basis of this eigendecomposition, we provide testable hypotheses pertaining to global properties of fly neuronal dynamics and highly specific circuits.

In the model we have proposed (Eq. 1), the eigenvectors of the effectome are patterns of neural activity which, if induced, will persist but scaled by the associated eigenvalue on each time step. For example if neural activity is set to the *i*th eigenvector (*x*_0_ = **v***_i_*) with eigenvalue *λ_i_*,

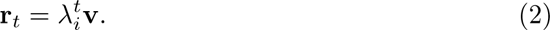

Thus the magnitude of the eigenvalue precisely determines the magnitude and duration of the effect of this pattern of activity. The eigenvectors with the largest eigenvalues are therefore plausibly associated with neural dynamics that have the largest total effects on the fly brain. The neurons associated with the significant coefficients, or ‘loadings’, in an eigenvector indicate the sub-population of neurons whose connectivity sustains these dynamics, forming an ‘eigencircuit’.

There are two critical properties of the eigendecomposition that determine the rate at which neural dynamics can be constrained by the estimated effectome. The first is the sparsity of eigenvectors. If the pattern of activity specified by the eigenvector includes only a handful of neurons with non-zero loadings, then only the effectome of those neurons needs to be learned to specify that dynamical mode. In contrast, if each eigenvector significantly involves all neurons, the entire effectome would need to be learned to explain even one dynamical mode. The second critical property is the relative magnitude of the eigenvalues. If the eigenvalue associated with a sparse eigenvector was far larger than all the others then the majority of variation in global dynamics could be explained with the effectome of a handful of neurons.

### Global dynamical properties of the putative effectome

We first examined the relative magnitude of the eigenvalues and found they decayed slowly. For example, the 1000th eigenvalue has 1*/*10 the effect as the largest (Fig 4A, black trace follows power-law *α* = *−*0.35 red trace). This implies that (1) the choice of which early modes to analyze is somewhat arbitrary because they have similar effect magnitudes on the brain, and (2) many dynamical modes are required to explain neural dynamics in the fly brain (i.e., fly neural dynamics are high-dimensional).

**Fig. 4:**
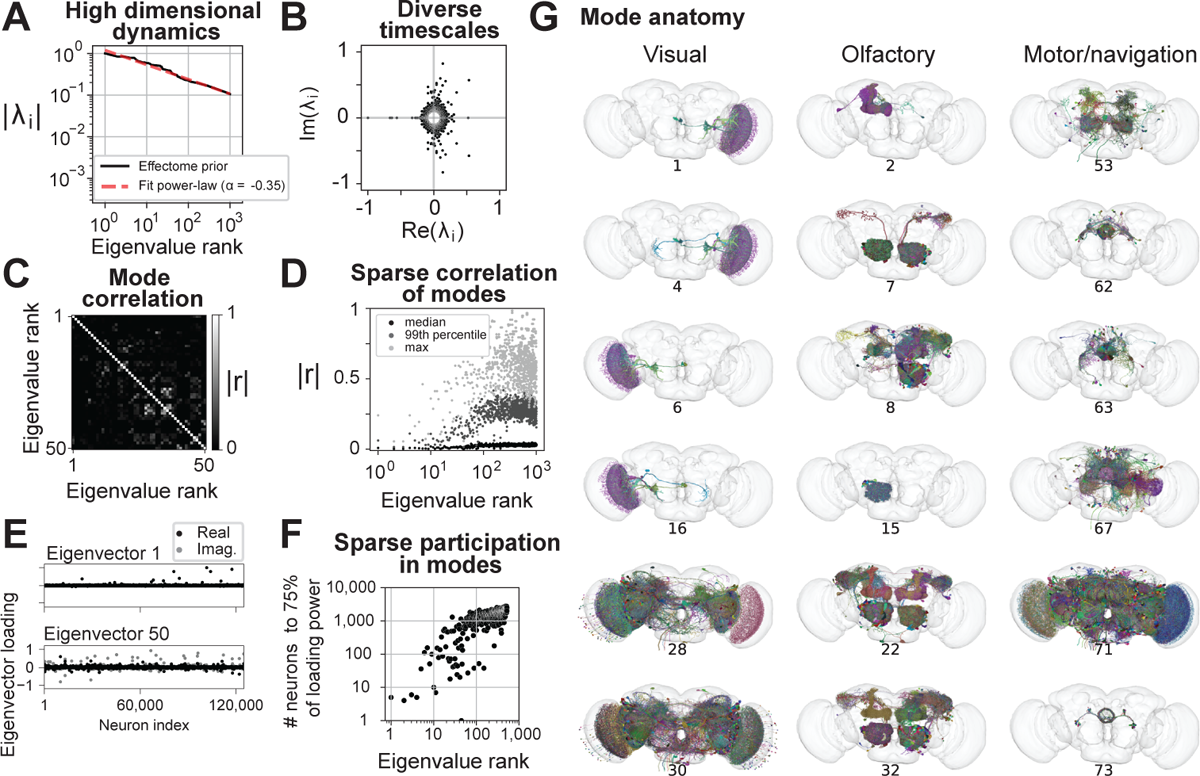
Putative global dynamical properties of the central fly nervous system. **(A)** Magnitude of top 1000 eigenvalues of the putative effectome (scaled matrix of signed synaptic counts extracted from the connectome, black) and power-law fit (dashed grey). **(B)** Eigen-values plotted in the complex plane. **(C)** Correlation of eigenvectors sorted by associated eigenvalue magnitude. **(D)** Median (black), 99th percentile (gray), and maximum (light grey) correlation between each eigenvector and all other top 1000 eigenvectors, showing that the first 10-20 eigenvectors are nearly orthogonal to other eigenvectors and the correlation between other eigenvectors is highly sparse. **(E)** Per-neuron eigenvector loadings for the first (top) and 50th (bottom) eigenvector. **(F)** Concentration of eigenvector loadings, quantified by the number of neurons needed to account for 75% of the eigen-vector’s power. **(G)** Anatomical renderings of neurons needed to account for 75% of the eigenvector’s loading power. Eigenvectors drawn from the top 100, and each column displays eigenvectors with neurons predominantly associated with either visual, olfactory, or motor/navigation anatomical locations.

We explored the range of time scales of predicted dynamics by examining the eigen-values in the complex plane (Fig 4B). Complex eigenvalues are associated with two orthogonal eigenvectors which define a plane of rotation. Intuitively, this implies that if a pattern of activity is induced that matches one of these eigenvectors, then over time it will transition into the second eigenvector. The angle from the positive real axis determines the speed of these rotational dynamics. At 0 degrees the eigenvalue is real positive and there are no oscillatory dynamics. A non-zero angle is exactly the angular step size of rotational dynamics at each time step, so small angles imply slow rotational dynamics. We find most eigenvalues are complex and distributed evenly with respect to angle; thus there is a broad distribution of time scales of rotational dynamics (scatter forms a circle). Very high magnitude eigenvalues tended to have large imaginary components reflecting rapid dynamics. The largest magnitude eigen-value was negative, implying the fastest possible rotational dynamics (flipping sign at each time step). Thus we predict the largest dynamical modes in the fly brain evolve at very rapid time scales.

We then investigated whether dynamical modes tended to be independent of each other. When eigenvectors are orthogonal to each other, the dynamics associated with each eigenvector are independent (under assumed white noise inputs). On the other hand, if two eigenvectors are highly correlated then these dynamics are more likely to co-occur. Thus by examining the correlation between eigenvectors we can ask which dynamical motifs will tend to be enlisted simultaneously—potentially because they are involved in similar computations. Examining the correlation matrix of the first 100 eigenvectors, we found on average correlation was weak with sparsely distributed higher values (Fig 4C). We noticed that early eigenvectors tended to have lower correlation to others (first 10 rows and columns dark). Aggregating the correlation of each of the top 1000 eigenvectors to all others, we found a clear trend where both the max and average increased with eigenvalue rank (Fig 4D). Roughly speaking, the top ten dynamical modes will occur independently of each other, while the rest will tend to co-occur with at least one other mode. Thus we predict dynamical modes will typically operate independently of each other, which bodes well for the project of examining these circuits individually.

In our analyses of how to feasibly fit the effectome experimentally, we determined that ideally sparse populations of neurons would be observed and stimulated. Thus here we analyzed whether the populations involved in these eigenmodes are in fact sparse. Plotting the first eigenvector we found that the loadings across neurons were extremely sparse, with the vast majority of loadings near zero and only several significantly deviating (Fig 4E top panel). These loading were of the same sign, indicating that all neurons significantly involved in this mode oscillate in sync. We found that later modes were also sparse but less so (bottom panel). To quantify sparsity across eigenvectors we measured the number of loadings needed to account for 75% of power across loadings (Fig 4F). For the first eigenvector only 5 neurons were needed (*<* 0.005% of all neurons). On average, for the top 10 eigenvectors *∼* 50 neurons were needed (*<* 0.05% of all neurons), for the 10th through 100th *∼* 500 neurons (*<* 0.5% of all neurons), and for the 100th through 1000th *∼* 1, 500 neurons (*<* 1.25% of all neurons). These findings suggest that the dynamics of the dominant modes in the fly brain can be explained by estimating a small fraction of the effectome.

Finally, we characterized the anatomical properties of these putative circuits. For each example circuit, we visualized the neurons that together comprise 75% of loading power within their respective eigenvector. To provide a broad sampling of circuits, we organized eigenvectors into three groups based on anatomical location: visual, olfactory, and motor/navigational (Figure 4G left, middle, and right columns respectively). In general, we found that the highly sparse top eigenvectors we previously characterized were also anatomically localized. For example, the top visual eigenvectors contained neurons confined to the lobula plate in the left hemisphere (rows 1-2) and right hemisphere (rows 3-4). These eigenvectors recapitulate a hypothesized neural circuit for opponent motion computation (Fig 4G, top left panel and see below). The top olfactory eigenvectors were also anatomically localized, and contained mushroom body neurons (row 1), projection neurons from the antennal lobe to the lateral horn (row 2), or local neurons in the antennal lobe (row 4). Multiple motor/navigational eigenvectors also showed confinement to neuropil regions including the LAL (row 1) and ellipsoid body (row 2, 3, 6). For all three anatomical categories, we observed that eigenvectors with lower sparsity tended to incorporate diverse cell types distributed across multiple neuropils (visual: rows 5,6; olfactory: rows 5,6; motor/navigational: rows 4,5). In general, we found that neurons with high loadings in early eigenvectors were often anatomically localized in accordance with the classic approach of studying the nervous system region by region. On the other hand some circuits were not anatomically localized and merit further investigation.

We discovered that the top eigenvectors of the connectome prior are highly sparse. This property facilitates learning the effectome because it is easier to identify, image, and stimulate sparse populations. Crucially, given the thousands of existing genetic driver lines accessible to the fly community [25] and tools for automatically screening these lines for neurons of interest [26], there is a high likelihood of identifying a reasonably sparse genetic line that contains the neurons of interest within any given eigencircuit (see Appendix B). Altogether, due to the distinct properties of the fly connectome—namely sparsity of connections, eigenvector loadings, and interaction between eigenmodes—there exists a plausible path forward to systematically and efficiently explain whole fly brain dynamics in terms of direct causal interactions between neurons.

### Putative effectome eigenvectors map onto interpretable circuits

Our decomposition of the putative effectome revealed sparse eigenvector loadings, which makes them amenable to further analysis. In particular, we asked whether these vectors correspond to identifiable circuits and interpretable dynamics in the fly brain (see Methods, ‘Simulations to analyze eigencircuits’).

We found that the first eigenvector localized to the lobula plate (Fig 5A) and was highly sparse (Fig 5B). The associated eigenvalue was real negative thus for linear dynamics we expect rapid oscillation, but in the more realistic case where activity is thresholded (see Methods) these neurons when activated will inactivate rapidly (Fig 5C). The top four neurons in this eigenvector were VCH, DCH, LPi2-1, and Am1 [3]. All putative effects between these neurons are inhibitory, but there is a complex mix of mutual and directed inhbition (Fig 5D, partial symmetry across diagonal of weight matrix). Notably VCH and DCH do not inhibit each other but are inhibited by Am1 and LPi2-1, but Am1 receives recurrent inhbition from VCAH and DCH while Lpi2-1 does not (Fig 5E).

**Fig. 5:**
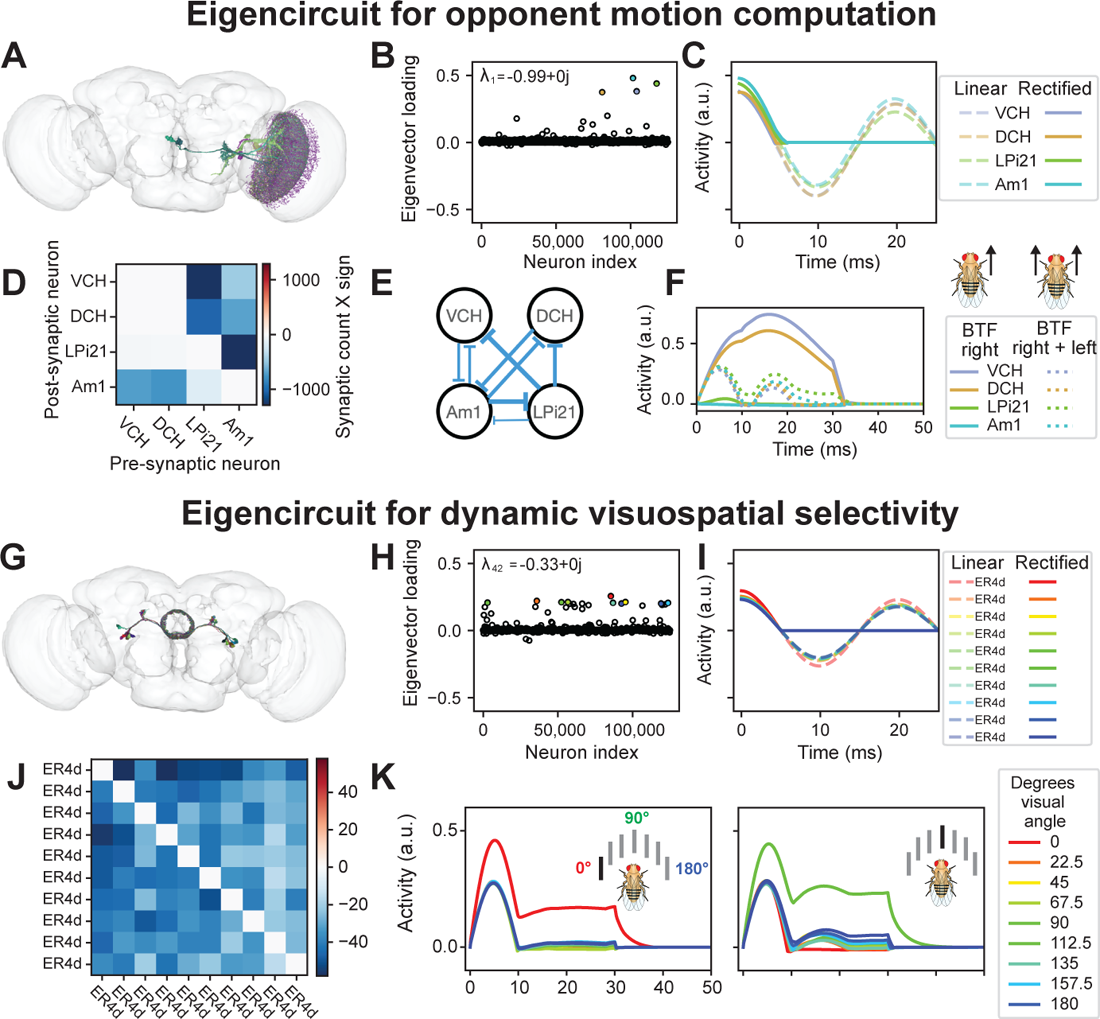
Example eigencircuits. **(A)** Neurons with top four loadings on eigenvector 1. **(B)** First eigenvector of fly effectome. **(C)** Linear and rectified dynamics upon stimulation by first eigenvector (dashed and solid traces). **(D)** Synaptic count and sign between neurons. **(E)** Circuit diagram representation of synaptic weight and sign. **(F)** Visual stimulation simulation. VCH and DCH are given sustained stimulation for 30 ms, simulating back-to-front (BTF) motion on the right side of the fly (left panel, solid orange and blue, high sustained response). All neurons were given sustained stimulation for 30 ms, simulating back-to-front (BTF) motion on both sides of the fly, which leads to inhibited responses of VCH and DCH (right panel, dashed orange and blue below solid). **(G)** Neurons with top four loadings on eigenvector 42. **(H)** All loadings for eigenvector 42. **(I)** Linear and rectified dynamics upon stimulation by eigenvector 42 (right panel). **(J)** Synaptic count and sign between 9 neurons with top loadings on eigenvector 42 (left panel). **(K)** Visual stimulation simulation. (left panel) simulation of a 0 degree visual stimulus, with strong input to the 0-preferring neuron (red trace) and sustained background input to all other neurons. The network response exhibits “winner take all” (WTA) dynamics, in which all neurons respond transiently to stimulus onset, but only the neuron with maximal response remains active for the entire stimulus duration. Right panel: similar results for a 90-degree stimulus and 90-degree preferring neuron (green trace).

Concretely determining the computation this visual circuit may perform requires an assumption about how visual features drive these neurons. It is known that VCH and DCH receive major input from H2 from the contralateral eye, which responds to back-to-front (BTF) motion [27], whereas, LPi2-1 and Am1 receive major inputs from T4b/T5b which are driven by ipsilateral BTF motion [28]. We then probed the functional properties of this circuit by simulating BTF visual input to either the right eye (contralateral input) or the left and right eyes together (bilateral input). When provided with contralateral BTF input only, we observe that VCH and DCH activity remain high throughout the stimulation period, while LPi2-1 and Am1 activity were suppressed because they were not directly stimulated and because Am1 is strongly inhibited by VCH and DCH (Fig 5F, solid trace blue and yellow above orange and green). On the other hand, bilateral BTF stimulation resulted in lower activity across the entire circuit, though only Am1 was fully suppressed during the stimulation period. Overall this circuit is well suited to compute opponent motion across the fly eyes: contralateral BTF motion activates DCH and VCH, which provide major inhibitory synaptic input to front-to-back selective T4a/T5a neurons in lobula plate layer 1.

Interestingly, this circuit was analyzed in a very recent small-scale connectomic analysis of the optic lobe [28] whereas it was ‘re-discovered’ with our data-driven 14 approach. This suggests, anecdotally, that the eigendecomposition of the connectome can reveal scientifically interesting sub-circuits.

Inspired by our findings with the first eigenvector of the connectome prior, we next sought to identify additional circuits with a putative role in stimulus selection. We found that eigenvector 42 contained high loadings for GABAergic R4d ring neurons in the ellipsoid body (Figure 5G), which have spatial receptive fields that retinotopically tile the visual field and exhibit directional and orientation tuning [29]. This eigenvector has sparse and bimodal loading (Fig 5H, majority of scatter near 0 but subset near 0.2). Its associated eigenvalue, similarly to the first eigenvector, is negative real leading to rapid oscillatory linear dynamics and inactivation for rectified dynamics (Fig 5I). Despite the similar dynamics, the synaptic weight matrix revealed complete mutual inhibition between R4d ring neurons (Fig 5J, off diagonal blue). This connectivity pattern has been identified in prior work [30], but its functional significance remains incompletely understood.

One possible computation consistent with this connectivity pattern is a winner-take-all (WTA) computation with respect to visual features distributed across space. In a WTA circuit, the most strongly activated neuron strongly suppresses all other neurons, thus preventing its own inhibition. To test this prediction, we simulated uniform visual drive to the R4d inhibitory sub-network while providing one neuron with higher input (Fig 5K left panel, stronger input to one neuron resulting in initial onset peak of red trace higher than that of others). We found that the neuron with a stronger input indeed had a robust sustained response while the other neurons response quickly decayed near to baseline (H, red trace at *∼* 0.25 after 10 ms but others near 0). We confirmed that this WTA property is not specific to a single neuron by providing different neurons with biased input (Fig 5K right panel). Indeed, these dynamics persisted, which supports the idea that a WTA computation is a robust property of this circuit.

Based on these findings, the R4d inhibitory sub-network appears well-posed to implement WTA dynamics and thus select individual visual-spatial channels as primary inputs to the central complex. The visual selectivity mechanism revealed through these simulations suggests a potential mechanism by which the central complex suppresses irrelevant or noisy visual inputs across space and hence maintains a stable heading representation.

We note that whereas global mutual inhibition within ring neurons such as the R4d cell type has been characterized [31] and predicted to drive WTA dynamics, to our knowledge we are the first to explicitly demonstrate that a mechanism predicted directly from anatomical parameters generates this computation.

We have analyzed two of the hundreds of sparse eigenvectors revealed by an eigendecomposition of the fly connectome. We emphasize that these eigenvectors should not be interpreted as the “true” effectome eigenvectors; rather, they provide a principled approach for generating and testing falsifiable hypotheses about causal relationships between neurons and the computations they may support. Our analyses served to demonstrate how neurons may be systematically chosen for causal perturbation experiments, and how—once the true effectome weights are learned for this subset of neurons—one might generate hypotheses about neural function than can be tested in-vivo. Crucially, the dynamical mechanisms of the computations indicated by our simulations remain untested, and require estimating the effectome between these neurons.

## Discussion

We developed a combined experimental and statistical approach to estimate a causal model of the fly central nervous system, its effectome. In simulation, we demonstrated that the approach provides consistent estimates of the ground truth effectome. We found the huge number of parameters of a whole brain model made this estimation infeasible, motivating us to use the connectome as a prior to drastically increase the efficiency of our estimator. We analyzed our ‘connectome prior’ to reveal thousands of small putative circuits operating largely independently of each other. This indicates that whole brain dynamics may be efficiently explained with sparse imaging and perturbation, which is far more feasible than dense imaging and perturbation. We analyzed two of these circuits to find that one recapitulates a proposed circuit for computing opponent motion and the other provides an explicit mechanism for visual spatial selectivity in the ellipsoid body.

### Synergies between the connectome and the effectome

The connectome clearly constrains how neurons can affect each other. Yet it is not clear exactly how. We believe a key motivation for accurately estimating causal effects is to build better models of how anatomy predicts physiology. In this way even partially learning the effectome can multiply the utility of the connectome. For example we explicitly assume in our ‘connectome prior’ that the magnitude of the effect between neurons is linearly proportional to the number of synapses, scaled by the same constant across the entire brain. By estimating the actual effects between neurons we can test and update this assumption. It is certainly plausible that individual synapses could have different weights leading to mis-matches between number of synapses and magnitude of effect. If this is found to be the case we could naturally encode it into our prior by placing a distribution on our hyper-parameters: for example, the synaptic scaling constant could be allowed to have some variation across neurons. Thus learning the effectome will not just provide causal circuit diagrams but increase the extent to which we can infer circuits from anatomy alone.

### Related work

Instrumental variables has been an area of intense interest outside of the neurosciences but was only recently recognized as a useful tool for neural perturbation analysis [11, 12]. Optogenetics has recently been employed as an instrumental variable on neuronal data [11]. Yet these approaches have focused on functional causal effects, they have not provided a method for separately identifying direct and indirect synaptic effects. The instrumental variables approach was recently extended to linear dynamical systems [32]. While this work does not consider an extension to the Bayesian setting nor how to design instrumental variables as we do here, they include extensions to the case of multiple lags at which observations effect each other (i.e., an AR(p) model with *p >* 1). This extension could be used to model different delays of synaptic effects and future work could place priors on these delays using axonal length and width.

### Linear approximation to nonlinear effects

The method we have introduced is a consistent estimator of linear effects between neurons yet the fly nervous system is a highly nonlinear system. One interpretation of the linear estimates in this setting are that they are a local approximation. For example they could accurately predict the effect of the hypothetical inclusion or exclusion of a single action potential but for a burst of many action potentials the accuracy would fall off.

The approach presented here could be extended to recurrent switching linear dynamical systems (rSLDS) [33].This class of models can approximate nonlinear dynamics similarly to how a set of line segments can approximate a curve.

### Technical limitations to learning the effectome

The success of our experimental framework hinges on (1) identifying genetic driver lines that target neurons of interest (i.e. neurons with high loadings in the connectome eigenvectors); (2) optogenetic stimulation with rank equal to number of neurons in the stimulated population (see Methods); (3) high temporal and spatial resolution recordings of neural dynamics; and (4) assigning connectome identities to each of the neurons that was imaged and optogenetically stimulated. The current spate of genetic driver lines and recently developed tools for screening these lines for neurons of interest makes (1) highly feasible. The remaining requirements do impose technological hurdles that must be addressed.

Although 2-photon holography has been used in other animal systems, most notably mice, to our knowledge no work has attempted to extend this technology to flies. This technological lag may be partly due to the fact that genetic drivers in the fly can result in exceptionally sparse labeling of neurons. As a result, probing specific neural circuits with optogenetics can be achieved through genetic, rather than optical, precision. However, we demonstrate in this work that both genetic and optical control are required to learn the effectome, thus necessitating the introduction of holography to the fly.

Our estimator also potentially requires rapid imaging speeds to uncover causal effects. In the most conservative case the time difference over which IV estimates are calculated would be the conduction delay of action potentials between the neurons being observed (1 ms conduction times typical [34]). Fortunately, the top neurons within each eigenvector are often anatomically localized, potentially meaning that only small sub-volumes of the brain need to be scanned during any given experiment. Nevertheless, the current acquisition speed available on most consumer 2-photon microscopes may still be insufficient to capture neural dynamics at the fastest timescales required to estimate a direct causal effect between neurons. Although custom microscopes have been engineered to achieve kilohertz-rate imaging [35], making this technology widely accessible to *Drosophila* neuroscientists will be a crucial step toward a community-driven effort to accurately estimate an effectome for the entire fly brain.

The last major technical hurdle toward estimating the effectome involves matching neurons that were recorded and perturbed experimentally with neurons in the connectome. Once again, the sparsity of each putative effectome circuit greatly facilitates this matching problem. Furthermore, recent work has successfully demonstrated registration of single cell types imaged at light level to the connectome [36]. However, whether existing registration approaches can unambiguously assign single neuron identities to each neuron in the experimental population remains untested. For a detailed strategy neuron assignment strategy, please refer to Appendix B.0.4.

## Methods

### Statistical model of fly brain

Here we approximate fly brain dynamics as a first order vector autoregressive model (VAR(1)),

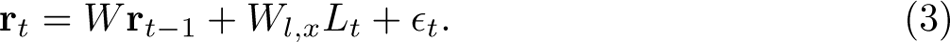

where **r***_t_* is the *D ×* 1 vector of responses from all neurons on time step *t*, *W* is the full *D × D* dynamics matrix, *L_t_* is the random *D_l_ ×* 1 vector of stimulation power (e.g., voltage), *W_l,x_* is the matrix specifying the effect of stimulation, and *ɛ_t_* is additive noise with arbitrary bounded covariance (for graphical model see supplementary figure S1). The raw instrumental variables estimator we outline below requires no distributional assumptions on the random variables (*L_t_*, *ɛ_t_*) but the derivation of the Bayesian-IV estimator requires normality.

### Simulations

#### Simulations to analyze eigencircuits

To evaluate neural dynamics associated with eigencircuits (see Results, ‘Putative effectome eigenvectors map onto interpretable circuits’) in continuous time we restricted our analyses to neurons with the highest eigenvector loadings that accounted for 75 % of all eigenvector power (for unit length eigenvectors equal to 1). In the case of the opponent motion circuit this was 5 neurons and in the case of the elipsoid body circuit this was 18 neurons. We set a time scale of *τ* = 10 ms and a sampling period of Δ = 1 ms so that at each time step,

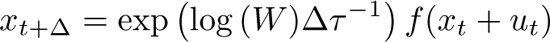

where *W* is the putative effectome of the subset of neurons being simulated, *f* (*·*) is a nonlinearity (here rectification), and *u_t_*is a vector time varying inputs.

#### Simulations to evaluate estimators

In all simulations to evaluate statistical estimators *ɛ_t_∼ N_D_*(0, Σ*_ɛ_*) where Σ*_ɛ_*= *I_D_* and *L_t_ ∼ N_D_*(0*, I_D_*). To simulate mis-specification of the connectome prior mean we estimated accuracy of our estimator across many ‘ground truth’ effectomes drawn from the connectome prior (except synaptic weights equal to 0 remained 0), such that the connectome prior was never the same as the ‘ground-truth’ effectome in a given simulation.

#### Instrumental variable estimator for a linear dynamical system

We will call the random vector of neurons that are observed in a given experiment, target neurons, at time point *t*, *Y_t_* and the neurons that are stimulated by the laser *X_t_*, source neurons (for graphical model see supplementary figure S1). In most cases we will assume that *Y_t_* includes the responses of the stimulated neurons as well. We then note that:

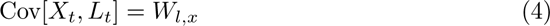

because the stimulation is assumed to have identity covariance. Thus by calculating the sample covariance between the laser and simultaneous activity in the stimulated neurons we can obtain an unbiased estimate of the linear weighting of laser drive on each neuron. Similarly we can obtain an unbiased estimate of the linear effect of the laser on all target neurons at the next time step,

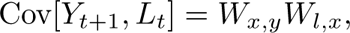

where *W_x,y_* is the submatrix of *W* with post-synaptic effects of *X_t_* on *Y_t_*_+1_. We can then use Eq. 4 to identify *W_x,y_*with,

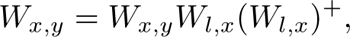

where (*W_l,x_*)^+^ is a pseudo inverse because we have not specified the rank of *L*. Only if *n_L_ ≥ n_S_* is this a true inverse and *W_l,x_* is invertible. An equivalent but more intuitive approach is termed two-stage least-squares(2SLS) where in the first stage *L_t_* is regressed on *X_t_* to give *X*^^^*_t_* = *W*^^^*_l,x_L_t_* and then *X*^^^*_t_* is regressed on *Y_t_*_+1_ to give *Y*^^^*_t_*_+1_= *W*^^^*_x,y_ X*^^^*_t_*. The IV estimator can also be extended to higher order AR processes [32].

#### Using the connectome as a prior for the parameters of the IV estimator

We use the 2SLS approach to motivate a consistent estimator from a Bayesian perspective. In short we perform classical Bayesian regression for the second stage of regression using the connectome as a prior on the weights *W_x,y_*. To be consistent with the most typical setting of Bayesian regression we first work out the case of multiple sources and a single target (learning a set of weights in the same row of *W*). We assume the conditional distribution of the output given the input is:

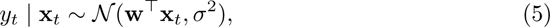

where (**x***_t_,* **y***_t_*) represents the input and output for sample *t ∈ {*1*, …, T}*, and *σ*^2^ is the variance of the observation noise in **y**.

Let us now suppose that *µ* provides the mean for a Gaussian prior over the linear weights **w**:

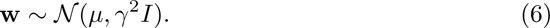

Let *µ* = *sc* where the hyperparameter *s* scales *c* the connectome prior we set to be equal to the synaptic count and sign. Combining this prior with the likelihood defined above gives us the following posterior mean:

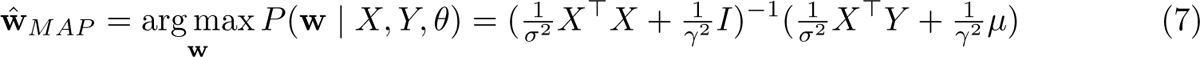

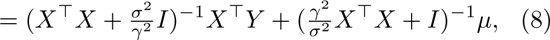

where *θ* = *{σ*^2^*, γ*^2^*, c}* denotes the hyperparameters. The second expression above (eq. 8) shows that the MAP estimate is the standard “ridge regression” estimate, **w**^ *ridge* = (*X^⊤^X* + *^σ^*^2^ *I*)*^−^*^1^*X^⊤^Y*, plus a term that biases the estimate towards the anatomical connectome *µ*. Note that in the limit of small observation noise *σ*^2^ or large prior variance *γ*^2^, the MAP estimate converges to the ML estimate, whereas in the limit of large *σ*^2^ or small *γ*^2^, it converges to *µ*.

In our simulations we choose the optimal hyperparameters beforehand but the hyperparameters could be learned via a standard cross-validation grid search. A more principled approach would be to use evidence optimization,

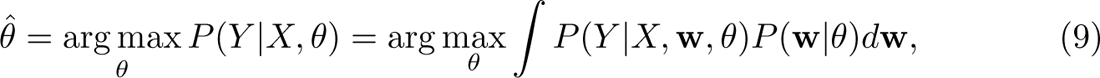

which would be straightforward given that the evidence is available in closed form for this model.

#### Number of experiments to identifiability

Here we propose to form a full effectome by performing experiments across flies. We analyse the identifiability of the effectome as a function of number of experiments. For identifiability of all post-synaptic weights for *n_S_* source neurons and a perturbation of rank 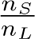, at least experiments are needed to identify the post-synaptic weights on *n_T_* target neurons. Thus to identify the weights on all *D* post-synaptic neurons will require repeating this procedure at least 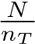 times. So overall the number of experiments needed to identify all of the source neurons post-synaptic weights (*n_E_*_(*S*)_) is,

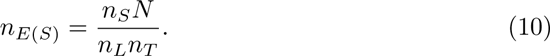

Then the total number of experiments (*N_E_*) to identify the entire effectome, assuming the number of source neurons is the same for each experiment is simply eq. 10 multiplied by 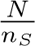,

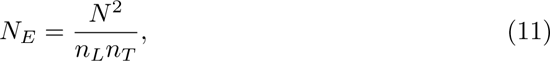

at worst this is *N* ^2^ and at best 1.

Identifiability does not guarantee an accurate estimate of the weights but is necessary for a consistent estimate. The amount of data needed to achieve an accurate estimate is a result of factors within single experiments, principally the SNR of the imaging modality, the strength of perturbation, and the number of samples. These factors are unknown in many of the experimental setups we recommend as they would vary from neuron-to-neuron, perturbation technique, and imaging technique. It seems plausible that within a single experiment enough samples could be collected to account for these other factors given recent approaches that allow up to 12 hours of continuous recording of neural activity in the fly [24].

## Acknowledgments

We thank Mala Murthy for feedback on the project and manuscript. We also thank Albert Lin for assistance with generating neuron renders. We thank Orren Tambour, Rich Pang, and Matt Creamer for insightful and inspiring discussions. We thank the Princeton FlyWire team and members of the Murthy and Seung labs, as well as members of the Allen Institute for Brain Science, for development and maintenance of FlyWire (supported by BRAIN Initiative grants MH117815 and NS126935 to Murthy and Seung). We acknowledge Jan Funke and his group for contributing neurotransmitter IDs. We also acknowledge members of the Princeton FlyWire team, led by Mala Murthy and Sebastian Seung, the Cambridge Drosophila Connectomics group led by Greg Jefferis, and the FlyWire consortium for neuron proofreading and annotation.

## Declarations

- Funding: JWP was supported by grants from the Simons Collaboration on the Global Brain (SCGB AWD543027) and the NIH BRAIN initiative (NS104899 and 9R01DA056404-04). DAP was supported Simons Collaboration on the Global Brain (SCGB AWD543027). MJA was supported by NIH NINDS R35 1R35NS111580-01 and BRAIN NINDS R01 NS104899.
- Availability of data and materials: all data is available at https://codex.flywire.ai/api/download
- Code availability: all code to analyze data and generate figures is available at https://github.com/dp4846/effectome

## Supplementary information

### Graphical model of IV estimator

The graphical model associated with our idealized experimental setup is depicted supplementary Figure S1. All variables are vectors and parameters are matrices. Dark arrows indicate estimands, grey arrows and variables respectively indicate parameters that cannot be estimated and variables that cannot be observed. *Z_t_*are unobserved neurons that potentially interact with observed populations. The most critical assumption is that optogenetic perturbation influences neural activity instantaneously (*L_t_* direct arrow in to *X_t_*) and do not directly affect other neurons or neural activity at other time points. In this model only neural activity in the previous time step directly effects neural activity in the next time step (arrows into *X_t_*, *Y_t_*, *Z_t_*only from *X_t−_*_1_, *Y_t−_*_1_, *Z_t−_*_1_). Note the similarity of the graph structure in Fig 1B and here: for example the linear relationship between *X_t−_*_1_ and *Y_t_* is the causal effect of interest and *Y_t−_*_1_ and *Z_t−_*_1_ are confounders. This approach can be extended to an AR(p) model to account for differing delays between neurons [32].

### Experimental approach to learning the effectome

An efficient strategy for estimating the fly effectome must consider multiple factors: (1) the total number of experiments to achieve full identifiability must be reasonably small

### Sparse expression

We propose expressing GCaMP and opsin in sparse subpopulations of neurons. Specifically, these neurons should correspond to those with the highest loadings in each of the eigenvectors obtained through eigendecomposition of the connectome prior (see Results, ‘Global dynamical properties of the putative effectome’). This prior can of course be updated as experiments progress. In the case where the neurons of interest have highly interdigitated neurites, segmenting these neurons for downstream identification may be infeasible. To address this, driver lines used may be further sparsened using the Sparse Predictive Activity through Recombinase Competition (SPARC) genetic toolkit [37]. SPARC will enable simultaneous expression of opsin and GCaMP in the same sparsened subpopulation of neurons, which is required by the approach we have outlined.

### Optogenetic activation

Estimation efficiency of the IV estimator increases as a function of effectively independent stimulation channels. To this end, we propose independently stimulating each labeled neuron in the brain using 2 photon holographic stimulation (HS). HS is a mature technology that can generate arbitrary light patterns in 3D volumes using spatial light modulators (SLMs). Multiple SLMs used in parallel can generate kilohertz-rate stimulation across approximately 1mm3 volumes [38], which is larger than the volume of the adult *Drosophila melanogaster* brain. Crucially, the sparse labeling outlined above will reduce the total laser power required to drive activity across the labeled population, thus preserving the health of the imaged fly.

### Registration to electron microscopy data

Creating the effectome for the fly brain requires mapping single-neuron dynamics to identified neurons. This is now possible with the whole-brain connectome [1] associated with the Full Adult Female Brain (FAFB) electron microscopy dataset [4]. To accomplish this, we propose following the BrIdge For Registering Over Statistical Templates (BIFROST) registration pipeline [36]. In short, the experimental fly will express a pan-neuronal structural marker (td-tomato) in addition to the opsin and GCaMP under control of the sparse genetic driver. This structure channel data (tomato-fast) will be acquired simultaneously with the functional GCaMP data. After the experiment is complete, a high-resolution scan of the GCaMP channel and td-tomato channel (tomato-slow) will be acquired. The high resolution scans acquired across experiments will be used to generate a mean brain template. Registration to the FAFB space is then accomplished through the following transformations: (a) tomato-fast to tomato-slow; (b) tomato-slow to mean brain; (c) mean brain to the standard functional *Drosophila* atlas (FDA); (d) FDA to FAFB. The first three steps rely on linear and nonlinear transformations, which can be accomplished using the ANTs library [39]. The final transformation is accomplished using a neural network called SynthMorph [40], which is able to bridge the disparate image statistics of the FDA and FAFB spaces. Finally, the transformations for the structural data will be applied to the functional GCaMP channel.

### Neural identification

The final step in our proposed experimental framework is to assign unique IDs to each neuron that expresses opsin and GCaMP. To do this, we suggest a two-pronged approach. First, the spatial footprint of ROIs extracted from the functional channel will be registered to the FAFB space, as outlined in (III). We expect that the ROIs may not perfectly align with the neural skeletons in the FAFB dataset. To address this, single-neuron light-level skeletons may be generated from the high-resolution anatomical scan obtained at the end of each experiment. NBLAST, an algorithm that computes anatomical similarities across datasets [41], may then be used to find candidate neurons in the FAFB datset with high anatomical similarity to the light-level neural skeleton that overlaps with a particular ROI. Critically, the location of the registered ROI in the FAFB space constrains the set of candidate neurons with which to perform NBLAST, thus greatly enhancing the efficiency of this process. The final neuron ID will then be determined by taking the consensus between the ROI registration approach and the top NBLAST candidates.

---

**Fig. 1:**
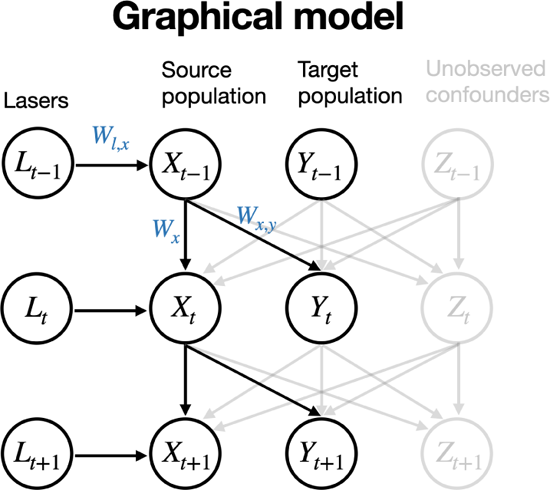
Graphical model associated with AR(1) estimation. *Lt* is IID stimulation at time step t, effect on source population is immediate and mediated by linear transformation *W_l,x_*. Effect of *X_t−_*_1_ on *Xt* and *Yt* (target, unstimulated population) is mediated respectively by linear transformation *Wx* and *Wx,y*. Interacting unobserved confounds can add arbitrary correlated noise (*Zt*).

(2) each imaged and stimulated neuron must be identifiable in the FAFB dataset; and
(3) total laser power exposed to the brain must be minimized to prevent heat-induced damage. To satisfy these requirements, we propose an experimental paradigm that relies on the following core components.

